# Nanoscale Organization of Membrane Tension during Neutrophil Extracellular Trap Formation Revealed by Fluorescence Lifetime Imaging

**DOI:** 10.64898/2026.04.29.718396

**Authors:** Jennifer Mohr, Linda Kartashew, Juliana Gretz, Sangeetha Shankar, Sebastian Jung, Sebastian Kruss, Luise Erpenbeck

**Author notes:** Shared first co-authors (Jennifer Mohr, Linda Kartashew, Juliana Gretz).

## Abstract

Cells continuously generate and respond to mechanical forces across compartments, with the plasma membrane acting as a nanoscale interface for sensing and transmitting tension. How intracellular forces are translated into membrane tension during dynamic processes such as neutrophil extracellular trap (NET) formation remains unclear. Here, we combine the mechanosensitive fluorescent probe Flipper-TR with fluorescence lifetime imaging microscopy (FLIM) to map spatiotemporal changes in plasma membrane tension in living cells. After validation in HeLa and dHL-60 cells under osmotic perturbation, we apply this approach to primary human neutrophils undergoing NETosis. Membrane tension transiently increases during chromatin decondensation and nuclear swelling within 90 min and is followed by a marked decrease 30 min after membrane rupture. Prior to rupture, tension is spatially heterogeneous, indicating localized nanoscale mechanical regulation. Cholesterol depletion abolishes the transient increase and reduces heterogeneity without affecting overall NETosis kinetics. Together, these findings establish the plasma membrane as a dynamic nanoscale reporter of intracellular mechanical stress during NETosis.

---

Cellular plasma membranes are dynamic lipid bilayers constantly exposed to mechanical forces during processes such as migration, phagocytosis, adhesion, and neutrophil extracellular trap (NET) formation. Membrane tension integrates inputs from lipid composition, membrane–cytoskeleton coupling, and membrane reservoirs (folds, microvilli, endomembrane stores) and thereby acts as a central regulator of cell morphology and function.^1–3^ However, how intracellular mechanical forces are translated into plasma membrane tension during highly dynamic cellular processes remains incompletely understood. This is particularly relevant for processes involving large-scale cellular remodeling, such as neutrophil extracellular trap (NET) formation.

Membrane tension influences a wide range of processes, including endocytosis, exocytosis, vesicle trafficking, and membrane remodeling, by constraining or facilitating vesicle budding and fusion and by modulating the recruitment and activity of curvature-generating proteins and trafficking machinery.^1–3^ Mechanical forces also propagate from the plasma membrane to the nucleus, where heterochromatin softening protects the genome against stress-induced damage.^4^ Effective membrane tension is closely linked to the local mechanical state of the bilayer, which reflects nanoscale lipid packing and organization rather than lipid composition per se. Under high tension, the membrane adopts a more ordered, less fluid state, whereas low tension favors increased molecular mobility and facilitates curvature generation, protein diffusion, and remodeling.^2,5^ These nanoscale changes occur below the diffraction limit and require sensitive biophysical readouts capable of resolving subtle variations in the local membrane environment.^2,5^ Classical approaches to quantify membrane tension include membrane tether pulling with optical tweezers or atomic force microscopy (AFM), where the steady-state tether force reports on effective membrane tension and membrane-to-cortex attachment. Other methods involve micropipette aspiration, membrane deformation by nanostructured substrates, and controlled osmotic perturbations to infer changes in tension and mechanical compliance.^5–7^ Although powerful, these methods are often low-throughput and may perturb the native state of the membrane, underscoring the need for minimally invasive, high-resolution mechanosensors or imaging.^5–7^

Environment-sensitive fluorescent probes have become key tools to visualize membrane mechanics in living cells with high spatiotemporal resolution.^3^ Flipper-TR is a landmark mechanosensitive dye that reports membrane tension through tension-dependent changes in its excited-state conformation and fluorescence lifetime.^8,9^ Incorporated into the lipid bilayer, Flipper-TR responds to alterations in lipid packing: increased tension and tighter packing stiffen its chromophore, leading to longer fluorescence lifetimes, whereas disordered, low-tension regions promote chromophore twisting and shorter lifetimes.^8,9^ Because fluorescence lifetime is largely independent of probe concentration, Flipper-TR enables quantitative mapping of nanoscale membrane order and tension in live cells.^8,9^ Such mechanosensitive probes, combined with advanced live-cell and *in vivo* imaging, provide a general framework to study how mechanical forces regulate cell behavior and tissue dynamics. Time-correlated single-photon counting (TCSPC) enables the measurement of fluorescence lifetimes and is not affected by intensity fluctuations due to photobleaching, scattering, or laser fluctuations.^10–12^ Fluorescence lifetime imaging microscopy (FLIM) combines laser scanning microscopy with TCSPC-based detection to generate spatial maps of fluorescence lifetimes.^13–15^ TCSPC-based FLIM represents a powerful modality to map membrane mechanics, probe Förster resonance energy transfer (FRET), resolve cellular metabolic states, and monitor protein–protein interactions, ion concentrations, and conformational changes in living cells.^12,14,16^

While the assessment of mechanical forces has predominantly focused on adherent cells and tissue remodeling, circulating immune cells are likewise exposed to defined mechanical constraints.^8,17^ In particular, neutrophils undergo pronounced mechanical transformations during activation, including intracellular pressure buildup and membrane tension changes associated with NET formation.

Neutrophilic granulocytes are fast-responding white blood cells that serve as a first line of defense against invading pathogens. ^18–21^ They are produced in the bone marrow, circulate in the bloodstream for up to 72 h, and, upon activation, deploy several antimicrobial strategies, including NET release and phagocytosis. ^18–24^ Excessive activation, as seen in sepsis or autoimmune disease, can lead to tissue damage. ^25,26^

NETosis is a distinct form of programmed cell death in which neutrophils release neutrophil extracellular traps (NETs) composed of decondensed DNA, histones, and granular and cytosolic proteins. ^23,24,27,28^ Building on our established biophysical model, ^29^ following activation, neutrophils engage enzymes such as neutrophil elastase (NE), myeloperoxidase (MPO), and PAD4, which promote chromatin decondensation, nuclear swelling, and nuclear envelope rupture. ^30–35^ This cascade increases intracellular pressure, drives cytoskeletal and endomembrane disassembly, and culminates in changes in plasma membrane tension and eventual rupture with NET release. ^33–36^

Notably, processes such as phagocytosis critically depend on membrane tension, lipid organization, and actin dynamics, and can be monitored using mechanosensitive probes such as Flipper-TR. However, membrane-interacting dyes may themselves affect such processes and should therefore be carefully controlled for.

Here, we use Flipper-TR combined with FLIM to quantify membrane tension in neutrophils during NETosis, revealing a characteristic spatiotemporal evolution of membrane tension consistent with pressure-driven buildup and subsequent relaxation, and uncovering pronounced nanoscale heterogeneity along the plasma membrane.

## Results

To investigate membrane mechanics in living cells, we employed the mechanosensitive fluorescent probe Flipper-TR, which reports changes in the local mechanical state of the plasma membrane through shifts in fluorescence lifetime. Figure 1A illustrates the structure of Flipper-TR and its expected orientation within the lipid bilayer under low- and high-tension conditions. Figure S1 depicts the experimental setup for time-correlated single-photon counting (TCSPC), and Figure 1B shows how conformational changes of Flipper-TR are translated into measurable fluorescence lifetimes. This framework enables quantitative, high-resolution mapping of nanoscale membrane mechanics across diverse cellular processes. We then applied this approach to neutrophils undergoing NET formation, a process characterized by pronounced mechanical remodelling.

**Figure 1.**
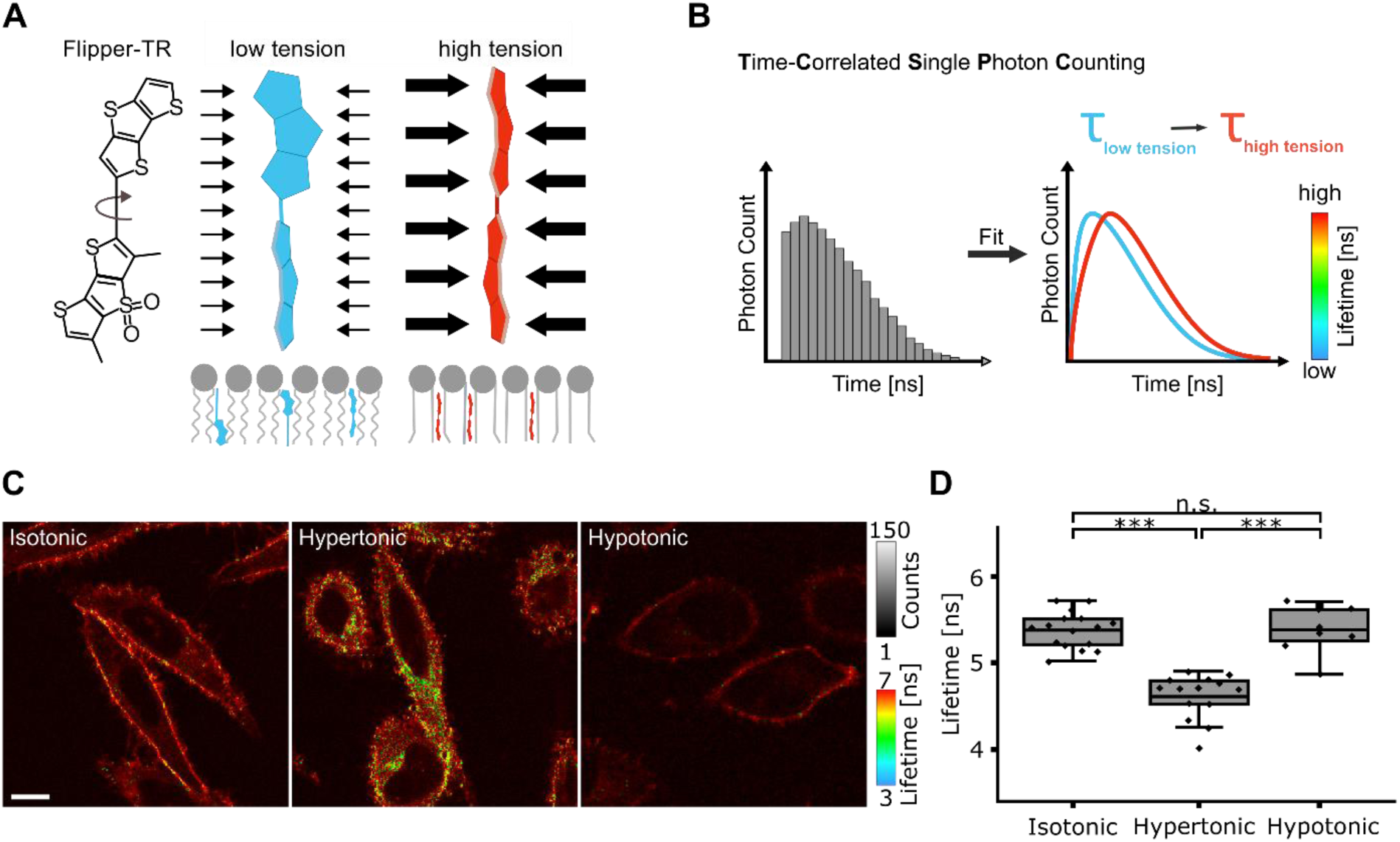
Fluorescence lifetime imaging of membrane tension in cells with the dye Flipper-TR. A) Structure of Flipper-TR and its expected orientation within a lipid bilayer under low- and high-tension conditions. B) Schematic of fluorescence lifetime measurement by Time-Correlated Single Photon Counting (TCSPC). C) HeLa cells exhibit differences lifetime under hypertonic conditions compared to isotonic conditions, but not under hypotonic conditions. (D) The mean lifetime of Flipper-TR in HeLa cells is reduced in hypertonic conditions (τ = 4.6 ± 0.3 ns) compared to isotonic (τ = 5.4 ± 0.3 ns) and hypotonic (τ = 5.4 ± 0.2 ns) conditions. (N ≤ 8 cells, box-and-whisker plots show median, interquartile range (box, 25–75th percentiles), and whiskers (5–95th percentiles), paired ANOVA: p-value *** <0.001, each data point corresponds to a single cell).

When exposed to different osmotic environments (1.6 M sucrose solution as hypertonic medium or ddH_2_O as hypotonic medium), HeLa cells exhibit characteristic changes in fluorescence lifetime when labelled with the mechanosensitive probe Flipper-TR. These lifetime variations report changes in the effective mechanical state of the membrane, which reflects membrane tension. Under isotonic and hypotonic conditions, the measured fluorescence lifetimes are comparable (τ= 5.4 ± 0.3 ns in an isotonic medium and τ= 5.4 ± 0.2 ns in a hypotonic medium) (Figure 1C, D). Hypotonic shock is expected to increase membrane tension due to water influx, which would typically lead to an increase in fluorescence lifetime. However, the observed lifetime shows little to no change. This suggests that cells effectively buffer acute increases in membrane tension, for example by recruiting membrane reservoirs or adjusting membrane–cytoskeleton coupling, thereby maintaining a relatively constant effective membrane tension under these conditions.

In contrast, hypertonic conditions (Figure 1C, middle) cause a marked decrease in fluorescence lifetime (τ = 4.6 ± 0.3 ns). This decrease is consistent with membrane relaxation and reduced membrane tension following water efflux. In this relaxed state, the observed lifetime decrease suggests a shift toward a more disordered membrane state rather than elevated membrane tension. While the underlying molecular changes are not directly resolved here, the data support a reduction in membrane order under hypertonic conditions.

This trend is not restricted to HeLa cells. Differentiated HL-60 cells, which are commonly used as a model for primary human neutrophils, display a similar behavior. Their fluorescence lifetime under hypertonic conditions (τ= 4.4 ns) is shorter than in an isotonic medium (τ= 5.3 ns), which is consistent with the mechanistic interpretation described above (Figure S2).

With Flipper-TR FLIM enabling quantitative mapping of membrane mechanics in living cells, we applied this approach to neutrophilic granulocytes to investigate membrane tension dynamics during NETosis, including in cholesterol-depleted cells (Figure 2A). To this end, we performed time-resolved fluorescence lifetime imaging of neutrophils undergoing PMA-induced NET formation stained with Flipper-TR. Intensity-based imaging of the plasma membrane (Flipper-TR) and nuclear staining (SPY505) enabled visualization of nuclear decondensation and chromatin expansion over time (Figure S3B).

**Figure 2.**
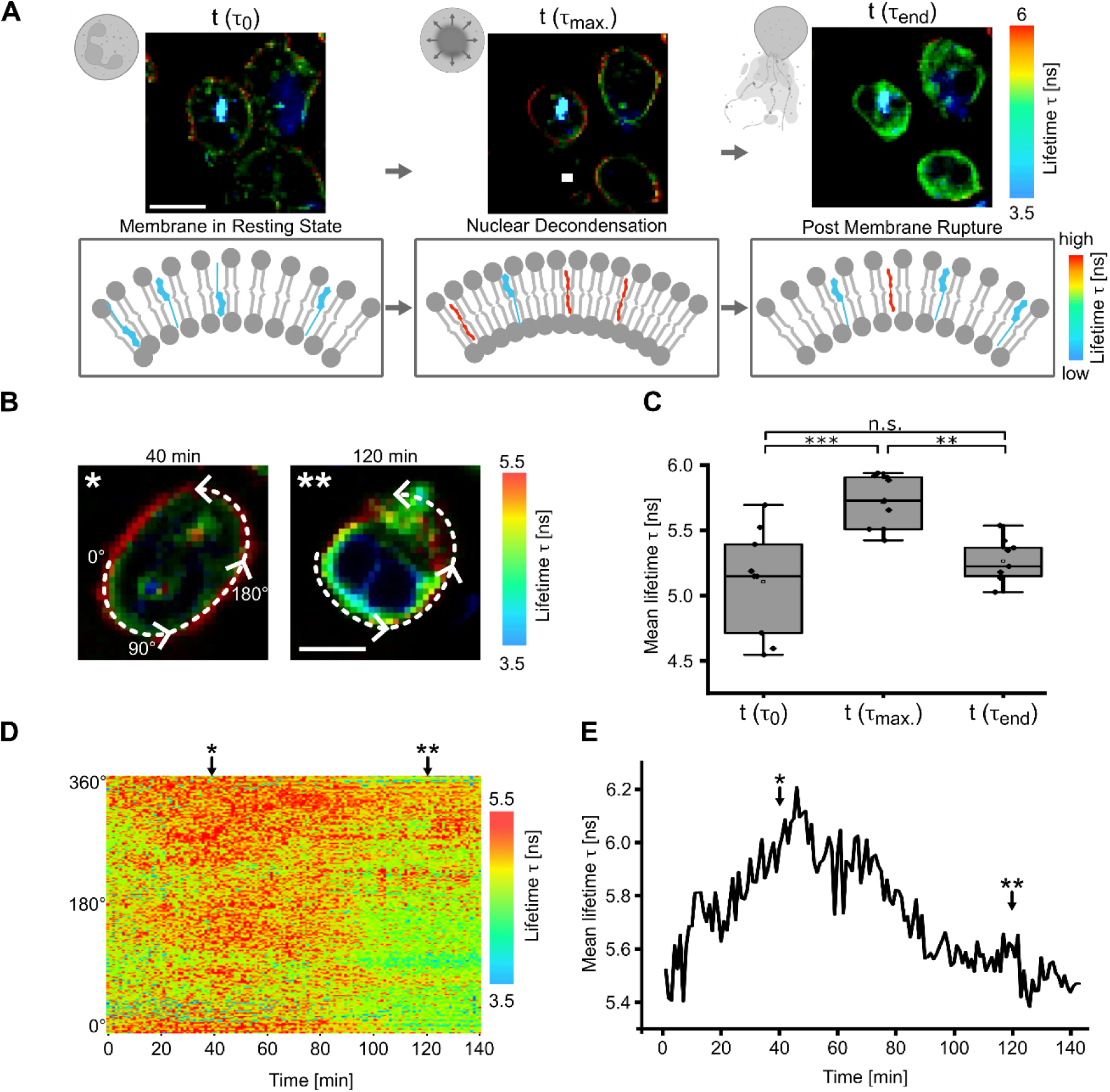
Spatiotemporal insights into membrane tension of neutrophilic granulocytes by fluorescence lifetime imaging. A) Temporal evolution of fluorescence lifetime (τ), reflecting membrane tension, during NETosis. Representative images at three time points show an initial increase in lifetime prior to membrane rupture, followed by a decrease 30 min after rupture. Upper schematics illustrate corresponding whole-cell morphology. Lower schematics depict the phospholipid bilayer with Flipper-TR integrated in its postulated orientation, sensing changes in membrane packing and tension. B) Methodology for lifetime (τ) data acquisition at specific time points. Each pixel along the membrane contour is sampled in a counter-clockwise manner, commencing from the left side of the cell. Exemplary data is shown for t = 40 minutes and t = 120 minutes. Scale bar accounts for 5 µm. C) Quantitative analysis of mean fluorescence lifetime (τ) at three distinct stages of NETosis: at the start, the point of membrane rupture, and 30 minutes post-rupture. Scale bar accounts for 10 µm. (n = 3 donors, N = 9 cells, box-and-whisker plots show median, interquartile range (box, 25–75th percentiles), and whiskers (5–95th percentiles), paired ANOVA: p-value ** < 0.01, *** <0.001, each data point corresponds to a single neutrophil). D) Heat map visualization of membrane lifetime (τ) values at each time point for the cell depicted in panel B. This representation provides a spatial and temporal overview of membrane tension dynamics. E) Mean fluorescence lifetime (τ) for the cell illustrated in panels B and D. The mean lifetime increases progressively until membrane rupture, followed by a gradual decline.

Analysis of fluorescence lifetime (τ) during NETosis revealed a characteristic temporal profile. In PMA-induced neutrophils, lifetime increased progressively following the onset of chromatin decondensation, peaked at the time of membrane rupture, and subsequently declined (Figure 2A). This increase is consistent with rising membrane tension driven by nuclear swelling, which promotes a more ordered membrane state and thereby prolongs fluorescence lifetimes. Quantification of mean lifetime values at defined stages— τ_0_ (imaging start), τ_max_ (membrane rupture), and τ_end_ (30 minutes post-rupture)—confirmed a significant transient increase in membrane tension prior to rupture, followed by a pronounced decrease (Figure 2C).

To resolve spatial heterogeneity in membrane tension, lifetime values were sampled along the plasma membrane contour in a counter-clockwise manner for each time point (Figure 2B). Heat map representations revealed that lifetime increases were not uniformly distributed but occurred in distinct, spatially confined regions of the membrane (Figure 2D and Figure S4A, S5A). These spatial variations likely reflect local differences in membrane organization, arising from region-specific mechanical constraints and heterogeneous coupling between the plasma membrane and the cytoskeleton. Consistently, such heterogeneity is compatible with non-uniform cytoskeletal disassembly and localized loss of membrane–cytoskeleton coupling during NETosis, potentially giving rise to region-specific mechanical instabilities.

Temporal analysis of mean lifetime values for individual cells further demonstrated a gradual increase in membrane tension prior to rupture, followed by a steady decline thereafter (Figure 2E, Figure S4A). Extending this framework, Flipper-TR FLIM revealed a characteristic tension trajectory that closely parallels the AFM-based tether force profile reported previously^29^: an initial increase in membrane tension during chromatin decondensation and nuclear swelling, followed by a decrease after plasma membrane rupture (Figure S6). The agreement between these fundamentally different measurement modalities supports that Flipper-TR lifetime faithfully captures NETosis-associated membrane tension dynamics, while providing massively increased spatiotemporal resolution along the plasma membrane.

Most cells completed NET formation within approximately 90 minutes after PMA stimulation, consistent with the observed kinetics of tension buildup. Following rupture, membrane tension reached a minimum within approximately 30 minutes, reflecting rapid loss of mechanical integrity. At this stage, the observed lifetime decrease is consistent with a shift toward a more disordered membrane state.

Together, these observations indicate that NETosis is accompanied by a dynamic and spatially heterogeneous redistribution of membrane tension. The progressive increase in lifetime during chromatin decondensation suggests that mechanical stress originating from nuclear expansion contributes to global membrane tension, while localized hotspots indicate additional region-specific regulation of membrane mechanics during NET formation.^29^

To assess how membrane material properties influence tension dynamics during NETosis, we analyzed neutrophils subjected to cholesterol depletion and compared them to untreated control cells. Cholesterol is a key regulator of plasma membrane mechanics. By intercalating between phospholipid acyl chains, it increases lipid packing density, modulates membrane fluidity, suppresses large-scale phase separation, and enhances the membrane’s ability to withstand and store mechanical stress.^37^

In neutrophils with intact cholesterol levels, these properties enable the plasma membrane to accumulate tension arising from intracellular forces during NETosis, including chromatin decondensation, nuclear expansion, and cytoskeletal remodeling, while maintaining a laterally heterogeneous but mechanically coupled membrane state. Previous studies have reported increased NETosis rates upon cholesterol depletion.^38,39^ In our experiments, all cells underwent NETosis following stimulation with 100 nM PMA, precluding direct assessment of changes in overall NETosis rates. However, tension dynamics (Figure 3C and Figure S4) allowed inference of NETosis onset based on the initial increase in membrane tension.

**Figure 3.**
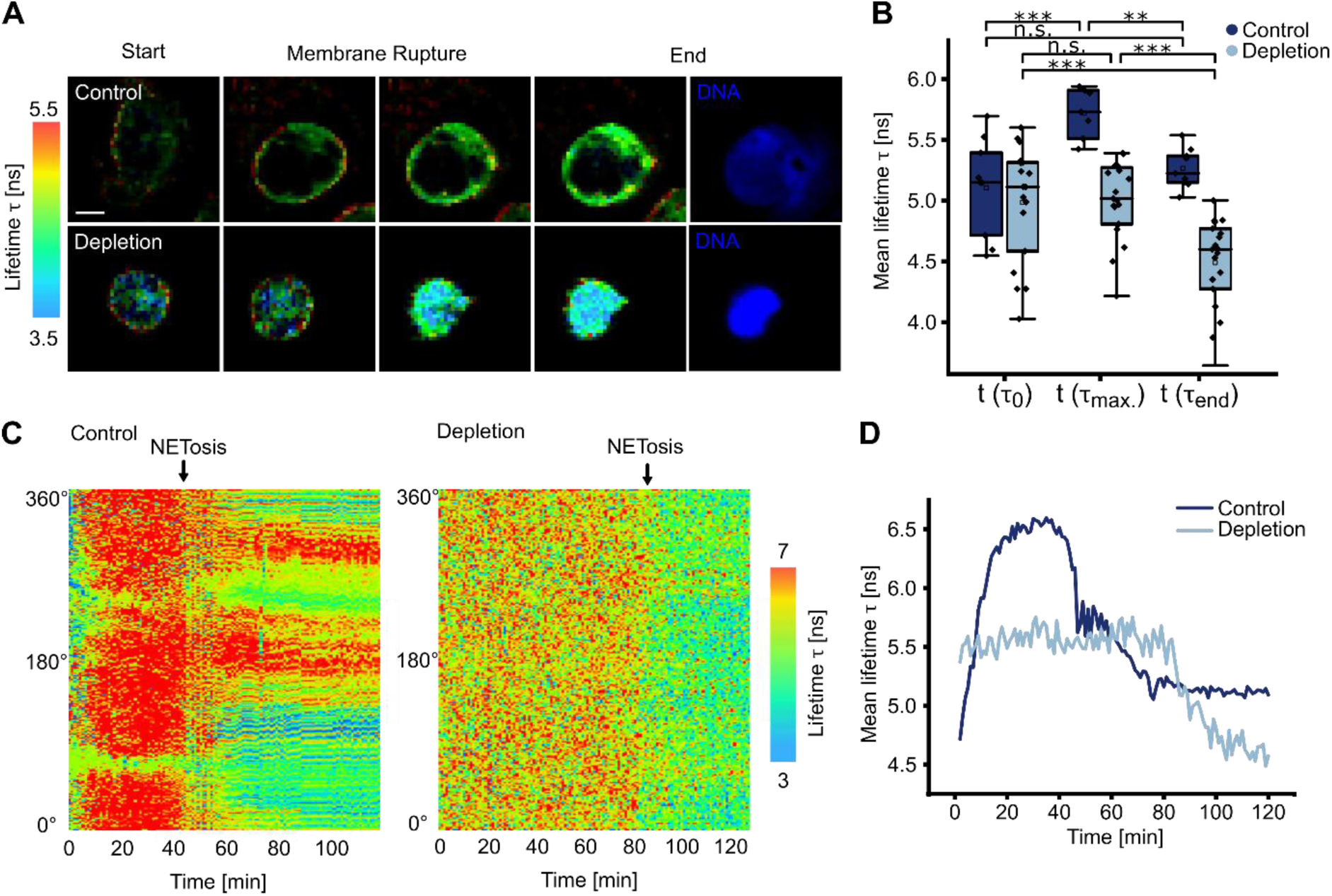
Spatiotemporal evolution of membrane tension in cholesterol depleted neutrophilic granulocytes during NETosis: A) Representative images of control and cholesterol-depleted neutrophils during PMA-induced NETosis at start of imaging, just before membrane rupture and 30 min post membrane rupture with a chromatin staining shown at the 30 min post-rupture time point. Cholesterol-depleted cells appear smaller and exhibit reduced expansion during NET formation compared to control cells. Scale bar: 10 µm. B) Temporal evolution of fluorescence lifetime (τ) as a proxy for membrane tension in control and cholesterol-depleted neutrophils. In control cells, τ increases from the start of imaging to the point of NETosis, followed by a decrease after membrane rupture. Cholesterol-depleted cells show no significant increase in τ during NETosis, reflecting impaired tension buildup, but both conditions display a comparable decrease post-rupture. (n ≤ 3 donors, N ≤ 9 cells box-and-whisker plots show median, interquartile range (box, 25–75th percentiles), and whiskers (5–95th percentiles), paired ANOVA: p-value ** < 0.01, *** <0.001, each data point corresponds to a single neutrophil). C) Heat map representation of spatiotemporal membrane tension distributions along the plasma membrane for control and cholesterol-depleted neutrophils. Control cells exhibit pronounced local maxima and minima, whereas cholesterol-depleted cells display a more homogeneous tension profile with reduced spatial variability. The point in time at which NETosis occurs is indicated by an arrow. D) Mean fluorescence lifetime (τ) for the cells illustrated in panels C. The mean lifetime increases progressively until membrane rupture in control cells, but shows no increase in the cholesterol-depleted cell, followed by a gradual decline after NETosis in both cells.

Cholesterol-depleted neutrophils exhibited distinct morphological differences, appearing smaller and displaying reduced cell expansion during NETosis compared to control cells (Figure 3A). Although based on two-dimensional imaging and not directly reporting volumetric changes, these observations are consistent with reduced membrane expansion.

Fluorescence lifetime analysis revealed marked differences in membrane tension between control and cholesterol-depleted neutrophils (Figure 3B). In control cells, membrane tension increased from the start of imaging to the point of membrane rupture and decreased thereafter. This transient is consistent with a shift towards a more ordered membrane state associated with tension buildup, likely reflecting increased lipid packing. In contrast, cholesterol-depleted neutrophils showed no significant change in lifetime prior to membrane rupture, indicating an impaired ability to build up the characteristic membrane mechanical state during chromatin decondensation and NET formation. This suggests that cholesterol contributes to efficient coupling between intracellular forces and membrane tension, whereas in its absence, mechanical stress is dissipated locally rather than accumulated globally. Both conditions exhibited a comparable decrease in lifetime after NETosis, consistent with loss of membrane integrity.

Spatial analysis further revealed that cholesterol depletion altered the distribution of membrane tension (Figure 3C,D, Figure S4). Control neutrophils displayed pronounced local maxima and minima, whereas cholesterol-depleted cells exhibited a more homogeneous tension profile with reduced spatial variability, also reflected in decreased variance (Figure 5). This indicates that cholesterol contributes to the formation of localized membrane domains capable of sustaining differential mechanical stress during NETosis.

Importantly, cholesterol depletion did not affect the overall kinetics of NETosis. The duration of PMA-induced NET formation was comparable between control and cholesterol-depleted neutrophils, with both conditions completing NETosis within approximately 90 minutes.

Together, these findings demonstrate that cholesterol is required for the characteristic buildup and spatial organization of the membrane mechanical state during NETosis, consistent with our established pressure-driven model, while overall NETosis kinetics remain unchanged.^29^

## Conclusion

This study establishes fluorescence lifetime imaging of the mechanosensitive probe Flipper-TR as a reliable reporter of membrane tension changes during immunological cell processes. In HeLa cells and differentiated HL-60 cells, Flipper-TR lifetimes responded predictably to osmotic perturbations, with reduced lifetimes under hypertonic conditions and no further increase under hypotonic compared to isotonic conditions, consistent with expected changes in membrane tension (Figure 1). Under hypotonic conditions, the lack of a further increase in lifetime despite elevated membrane tension suggests that the increase in effective tension may be counterbalanced by concomitant changes in membrane organization, such as enhanced lateral heterogeneity, which could dampen the net effect on the observed lifetime. These responses reflect the combined effects of osmotic water flux and changes in the membrane mechanical state, with opposing biophysical contributions under hypotonic conditions effectively compensating each other. Together, these experiments validate Flipper-TR lifetime as a quantitative readout of the membrane mechanical state and complement previous measurements of neutrophil membrane tension by atomic force microscopy.^29^ While our validation experiments support Flipper-TR lifetime as a quantitative readout of the membrane mechanical state, it should be considered that membrane-intercalating dyes are not inherently inert and therefore require appropriate biological controls.

Applying this approach to neutrophils undergoing PMA-induced NETosis revealed a transient increase in membrane tension preceding membrane rupture, followed by a decrease after loss of membrane integrity. The increase in fluorescence lifetime during chromatin decondensation is consistent with a transient shift toward a more ordered membrane state, whereas the post-rupture decrease reflects membrane relaxation and loss of mechanical coupling. Spatially resolved lifetime analysis further revealed heterogeneous membrane tension distributions, reflecting nanoscale organization of membrane mechanical states (Figure 2, Figure S4A). Control cells exhibited pronounced spatial variance with distinct local maxima and minima.

These observations are consistent with a biophysical view of NETosis in which intracellular mechanical remodeling contributes to membrane deformation and rupture. Previous work has identified chromatin swelling as a major contributor to intracellular force generation during NET formation.^29^ Our data extend this concept by demonstrating that these forces are not reflected in a uniform membrane response, but instead give rise to a spatially heterogeneous membrane mechanical state. This suggests that membrane mechanics during NETosis is organized at the nanoscale rather than being globally homogeneous.

Consistently, cholesterol depletion altered both the temporal evolution and spatial organization of membrane tension. Cholesterol-depleted cells showed no increase in membrane tension prior to rupture and displayed a more homogeneous membrane tension profile with reduced spatial variability (Figure 3, Figure S4B). This indicates that membrane material properties, and in particular cholesterol-dependent lipid organization, are required for the buildup and spatial patterning of membrane tension during NETosis. Notably, overall NETosis kinetics remained unchanged, suggesting that membrane composition primarily affects the nanoscale organization of the membrane mechanical state rather than the overall progression of NET formation.

More broadly, these findings position membrane tension as part of a mechanically regulated layer of NETosis that complements established biochemical pathways. In addition to cytoskeletal remodeling and intracellular force generation, membrane material properties contribute to how mechanical stresses are distributed and resolved at the plasma membrane. The ability to resolve spatial and temporal variations in membrane tension provides direct experimental access to the nanoscale organization of membrane mechanical states during dynamic cellular processes.

## AUTHOR INFORMATION

### Author Contributions

SK and LE conceived the project. LK implemented the methodology. LK and JM performed the investigation. LK, JM, JG and SS were responsible for data visualization. SK and LE acquired funding, administered the project, and supervised the research. JM and SK wrote the original draft of the manuscript. JM, JG, SK, SJ, and LE viewed and edited the manuscript. All authors have given approval to the final version of the manuscript. ‡These authors contributed equally.

### Diversity, equity, ethics, and inclusion

This study was approved by the Ethics Committee Westfalen-Lippe (approval number 2021-657-f-S). Before donating blood, fully informed consent of each donor was obtained.

## ACKNOWLEDGMENT

We thank all blood donors for their valuable contribution and everyone who helped with blood sample collection. We thank the Fonds der chemischen Industrie (FCI) for funding to improve teaching. We thank the DFG for funding.

## SUPPORTING INFORMATION

Experimental setup; membrane tension changes under hypo- and hypertonic conditions in differentiated HL-60 cells; chromatin staining during NETosis and TCSPC false-color imaging; heat map visualization of membrane lifetime *τ* values at each time point for exemplary cells; variance of lifetime values for the cells shown in Figures 2D and 2E; comparison of lifetime data with AFM measurements during NETosis (PDF).

## REFERENCES

(1) Gauthier, N. C.; Masters, T. A.; Sheetz, M. P. Mechanical Feedback between Membrane Tension and Dynamics. Trends Cell Biol. 2012, 22 (10), 527–535. 10.1016/J.TCB.2012.07.005.

(2) Yan, Q.; Perez, C. G.; Karatekin, E. Cell Membrane Tension Gradients, Membrane Flows, and Cellular Processes. Physiology 2024, 39 (4), 231. 10.1152/PHYSIOL.00007.2024.

(3) Sitarska, E.; Diz-Muñoz, A. Pay Attention to Membrane Tension: Mechanobiology of the Cell Surface. Curr. Opin. Cell Biol. 2020, 66, 11. 10.1016/J.CEB.2020.04.001.

(4) Nava, M. M.; Miroshnikova, Y. A.; Biggs, L. C.; Whitefield, D. B.; Metge, F.; Boucas, J.; Vihinen, H.; Jokitalo, E.; Li, X.; García Arcos, J. M.; Hoffmann, B.; Merkel, R.; Niessen, C. M.; Dahl, K. N.; Wickström, S. A. Heterochromatin-Driven Nuclear Softening Protects the Genome against Mechanical Stress-Induced Damage. Cell 2020, 181 (4), 800–817.e22. 10.1016/j.cell.2020.03.052.

(5) Roy, D.; Steinkühler, J.; Zhao, Z.; Lipowsky, R.; Dimova, R. Mechanical Tension of Biomembranes Can Be Measured by Super Resolution (STED) Microscopy of Force-Induced Nanotubes. Nano Lett. 2020, 20 (5), 3185–3191. 10.1021/ACS.NANOLETT.9B05232/SUPPL_FILE/NL9B05232_SI_002.AVI.

(6) Lüchtefeld, I.; Zurich, E.; Zurich, S. Studying Cell Membrane Tension and Mechanosensation during Mechanical Stimulation. Nature Methods 2024 21:6 2024, 21 (6), 944–945. 10.1038/s41592-024-02278-7.

(7) Kreysing, E.; Hugh, J. M.; Foster, S. K.; Andresen, K.; Greenhalgh, R. D.; Pillai, E. K.; Dimitracopoulos, A.; Keyser, U. F.; Franze, K. Effective Cell Membrane Tension Is Independent of Polyacrylamide Substrate Stiffness. PNAS Nexus 2023, 2 (1), 1–8. 10.1093/PNASNEXUS/PGAC299.

(8) Colom, A.; Derivery, E.; Soleimanpour, S.; Tomba, C.; Molin, M. D.; Sakai, N.; González-Gaitán, M.; Matile, S.; Roux, A. A Fluorescent Membrane Tension Probe. Nat. Chem. 2018, 10 (11), 1118–1125. 10.1038/S41557-018-0127-3.

(9) Dal Molin, M.; Verolet, Q.; Colom, A.; Letrun, R.; Derivery, E.; Gonzalez-Gaitan, M.; Vauthey, E.; Roux, A.; Sakai, N.; Matile, S. Fluorescent Flippers for Mechanosensitive Membrane Probes. J. Am. Chem. Soc. 2015, 137 (2), 568–571. 10.1021/JA5107018.

(10) Bitton, A.; Sambrano, J.; Valentino, S.; Houston, J. P. A Review of New High-Throughput Methods Designed for Fluorescence Lifetime Sensing From Cells and Tissues. Front. Phys. 2021, 9, 648553. 10.3389/FPHY.2021.648553.

(11) Spatola Rossi, T.; Pain, C.; Botchway, S. W.; Kriechbaumer, V. FRET-FLIM to Determine Protein Interactions and Membrane Topology of Enzyme Complexes. Curr. Protoc. 2022, 2 (10), e598. 10.1002/CPZ1.598.

(12) Trinh, A. L.; Esposito, A. Biochemical Resolving Power of Fluorescence Lifetime Imaging: Untangling the Roles of the Instrument Response Function and Photon-Statistics. Biomed. Opt. Express 2021, 12 (7), 3775. 10.1364/BOE.428070.

(13) Duncan, R. R.; Bergmann, A.; Cousin, M. A.; Apps, D. K.; Shipston, M. J. Multi-Dimensional Time-Correlated Single Photon Counting (TCSPC) Fluorescence Lifetime Imaging Microscopy (FLIM) to Detect FRET in Cells. J. Microsc. 2004, 215 (Pt 1), 1. 10.1111/J.0022-2720.2004.01343.X.

(14) Becker, W.; Bergmann, A.; Hink, M. A.; König, K.; Benndorf, K.; Biskup, C. Fluorescence Lifetime Imaging by Time-Correlated Single-Photon Counting. Microsc. Res. Tech. 2004, 63 (1), 58–66. 10.1002/JEMT.10421.

(15) Kumar, V.; Schlücker, S.; Hasselbrink, E. Ultrafast Time-Resolved Molecular Spectroscopy. Molecular and Laser Spectroscopy: Advances and Applications: Volume 2 2020, 563–594. 10.1016/B978-0-12-818870-5.00016-2.

(16) Bitton, A.; Sambrano, J.; Valentino, S.; Houston, J. P. A Review of New High-Throughput Methods Designed for Fluorescence Lifetime Sensing From Cells and Tissues. Front. Phys. 2021, 9, 648553. 10.3389/FPHY.2021.648553.

(17) García-Arcos, J. M.; Mehidi, A.; Sanchez-Velasquez, J.; Guillamat, P.; Tomba, C.; Houzet, L.; Capolupo, L.; Espadas, J.; D’Angelo, G.; Colom, A.; Hinde, E.; Aumeier, C.; Roux, A. Adherent Cells Sustain Membrane Tension Gradients Independently of Migration. Nat. Commun. 2025, 16 (1), 10539. 10.1038/s41467-025-65571-9.

(18) Bartneck, M.; Wang, J. Therapeutic Targeting of Neutrophil Granulocytes in Inflammatory Liver Disease. Front. Immunol. 2019, 10. 10.3389/FIMMU.2019.02257.

(19) Zheng, Y.; Sefik, E.; Astle, J.; Karatepe, K.; Öz, H. H.; Solis, A. G.; Jackson, R.; Luo, H. R.; Bruscia, E. M.; Halene, S.; Shan, L.;; Flavell, R. A. Human Neutrophil Development and Functionality Are Enabled in a Humanized Mouse Model. Proceedings of the National Academy of Sciences (PNAS) 2022. 10.1073/pnas.

(20) Silvestre-Roig, C.; Fridlender, Z. G.; Glogauer, M.; Scapini, P. Neutrophil Diversity in Health and Disease. Trends Immunol. 2019, 40 (7), 565. 10.1016/J.IT.2019.04.012.

(21) Koenderman, L.; Tesselaar, K.; Vrisekoop, N. Human Neutrophil Kinetics: A Call to Revisit Old Evidence. Trends Immunol. 2022, 43 (11), 868–876. 10.1016/J.IT.2022.09.008.

(22) Uribe-Querol, E.; Rosales, C. Phagocytosis: Our Current Understanding of a Universal Biological Process. Front. Immunol. 2020, 11, 1066. 10.3389/FIMMU.2020.01066.

(23) Zeng, W.; Song, Y.; Wang, R.; He, R.; Wang, T. Neutrophil Elastase: From Mechanisms to Therapeutic Potential. J. Pharm. Anal. 2023, 13 (4), 355. 10.1016/J.JPHA.2022.12.003.

(24) Monteith, A. J.; Miller, J. M.; Maxwell, C. N.; Chazin, W. J.; Skaar, E. P. Neutrophil Extracellular Traps Enhance Macrophage Killing of Bacterial Pathogens. Sci. Adv. 2021, 7 (37). 10.1126/SCIADV.ABJ2101/SUPPL_FILE/SCIADV.ABJ2101_SM.PDF.

(25) McKenna, E.; Mhaonaigh, A. U.; Wubben, R.; Dwivedi, A.; Hurley, T.; Kelly, L. A.; Stevenson, N. J.; Little, M. A.; Molloy, E. J. Neutrophils: Need for Standardized Nomenclature. Front. Immunol. 2021, 12, 602963. 10.3389/FIMMU.2021.602963/XML/NLM.

(26) Rosales, C. Neutrophil: A Cell with Many Roles in Inflammation or Several Cell Types? Front. Physiol. 2018, 9 (FEB), 113. 10.3389/FPHYS.2018.00113.

(27) Petretto, A.; Bruschi, M.; Pratesi, F.; Croia, C.; Candiano, G.; Ghiggeri, G.; Migliorini, P. Neutrophil Extracellular Traps (NET) Induced by Different Stimuli: A Comparative Proteomic Analysis. PLoS One 2019, 14 (7), e0218946. 10.1371/JOURNAL.PONE.0218946.

(28) Pires, R. H.; Felix, S. B.; Delcea, M. The Architecture of Neutrophil Extracellular Traps Investigated by Atomic Force Microscopy. Nanoscale 2016, 8 (29), 14193–14202. 10.1039/C6NR03416K.

(29) Neubert, E.; Meyer, D.; Rocca, F.; Günay, G.; Kwaczala-Tessmann, A.; Grandke, J.; Senger-Sander, S.; Geisler, C.; Egner, A.; Schön, M. P.; Erpenbeck, L.; Kruss, S. Chromatin Swelling Drives Neutrophil Extracellular Trap Release. Nature Communications 2018 9:1 2018, 9 (1), 3767-. 10.1038/s41467-018-06263-5.

(30) Kearney, P. L.; Bhatia, M.; Jones, N. G.; Yuan, L.; Glascock, M. C.; Catchings, K. L.; Yamada, M.; Thompson, P. R. Kinetic Characterization of Protein Arginine Deiminase 4: A Transcriptional Corepressor Implicated in the Onset and Progression of Rheumatoid Arthritis. Biochemistry 2005, 44 (31), 10570–10582. 10.1021/BI050292M/ASSET/IMAGES/LARGE/BI050292MF00008.JPEG.

(31) Metzler, K. D.; Fuchs, T. A.; Nauseef, W. M.; Reumaux, D.; Roesler, J.; Schulze, I.; Wahn, V.; Papayannopoulos, V.; Zychlinsky, A. Myeloperoxidase Is Required for Neutrophil Extracellular Trap Formation: Implications for Innate Immunity. Blood 2011, 117 (3), 953–959. 10.1182/BLOOD-2010-06-290171.

(32) Papayannopoulos, V.; Metzler, K. D.; Hakkim, A.; Zychlinsky, A. Neutrophil Elastase and Myeloperoxidase Regulate the Formation of Neutrophil Extracellular Traps. Journal of Cell Biology 2010, 191 (3), 677–691. 10.1083/JCB.201006052.

(33) Gößwein, S.; Lindemann, A.; Mahajan, A.; Maueröder, C.; Martini, E.; Patankar, J.; Schett, G.; Becker, C.; Wirtz, S.; Naumann-Bartsch, N.; Bianchi, M. E.; Greer, P. A.; Lochnit, G.; Herrmann, M.; Neurath, M. F.; Leppkes, M. Citrullination Licenses Calpain to Decondense Nuclei in Neutrophil Extracellular Trap Formation. Front. Immunol. 2019, 10 (OCT), 485362. 10.3389/FIMMU.2019.02481/BIBTEX.

(34) Thiama, H. R.; Wong, S. L.; Qiu, R.; Kittisopikul, M.; Vahabikashi, A.; Goldman, A. E.; Goldman, R. D.; Wagner, D. D.; Waterman, C. M. NETosis Proceeds by Cytoskeleton and Endomembrane Disassembly and PAD4-Mediated Chromatin Decondensation and Nuclear Envelope Rupture. Proceedings of the National Academy of Sciences (PNAS) 2020, 117 (13), 7326–7337. 10.1073/PNAS.1909546117/SUPPL_FILE/PNAS.1909546117.SM22.AVI.

(35) Li, Y.; Li, M.; Weigel, B.; Mall, M.; Werth, V. P.; Liu, M. Nuclear Envelope Rupture and NET Formation Is Driven by PKCα-mediated Lamin B Disassembly. EMBO Rep. 2020, 21 (8). 10.15252/EMBR.201948779/SUPPL_FILE/EMBR201948779-SUP-0008-SDATAEVFIGS.ZIP.

(36) Thiam, H. R.; Wong, S. L.; Wagner, D. D.; Waterman, C. M. Cellular Mechanisms of NETosis. Annu. Rev. Cell Dev. Biol. 2020, 36, 191. 10.1146/ANNUREV-CELLBIO-020520-111016.

(37) Mangiarotti, A.; Sabri, E.; Schmidt, K. V.; Hoffmann, C.; Milovanovic, D.; Lipowsky, R.; Dimova, R. Lipid Packing and Cholesterol Content Regulate Membrane Wetting and Remodeling by Biomolecular Condensates. Nat. Commun. 2025, 16 (1), 2756. 10.1038/S41467-025-57985-2.

(38) Neumann, A.; Brogden, G.; Jerjomiceva, N.; Brodesser, S.; Naim, H. Y.; Von Köckritz-Blickwede, M. Lipid Alterations in Human Blood-Derived Neutrophils Lead to Formation of Neutrophil Extracellular Traps. Eur. J. Cell Biol. 2014, 93 (8–9), 347–354. 10.1016/J.EJCB.2014.07.005.

(39) Henneck, T.; Mergani, A.; Clever, S.; Seidler, A. E.; Brogden, G.; Runft, S.; Baumgärtner, W.; Branitzki-Heinemann, K.; von Köckritz-Blickwede, M. V. Formation of Neutrophil Extracellular Traps by Reduction of Cellular Cholesterol Is Independent of Oxygen and HIF-1α. Int. J. Mol. Sci. 2022, 23 (6), 3195. 10.3390/IJMS23063195/S1.

