## Supplementary Information for "Nanoscale Organization of Membrane Tension during Neutrophil Extracellular Trap Formation Revealed by Fluorescence Lifetime Imaging"

‡ Shared first co-authors

* Corresponding authors

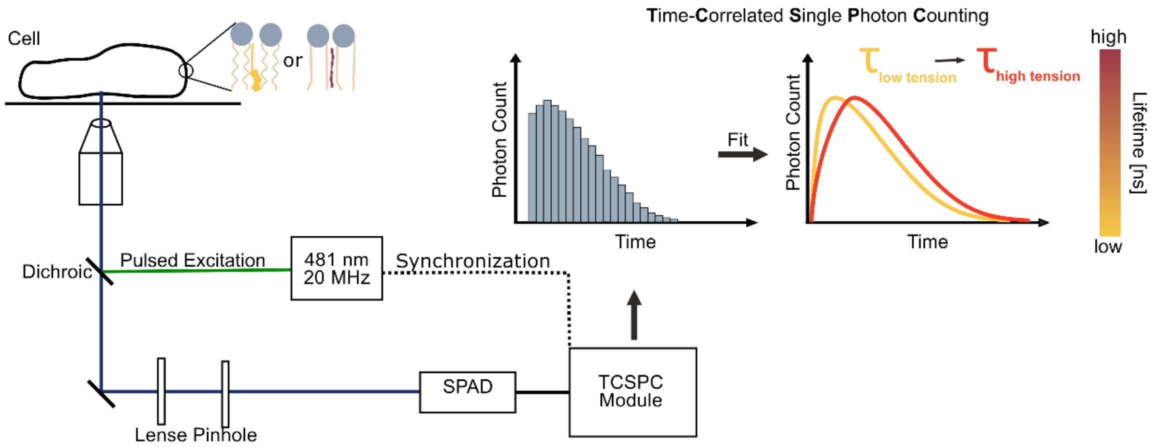


**Figure S1**. Experimental setup using Time-Correlated Single Photon Counting (TCSPC) and the approach to use lifetime measurements to access membrane tension.

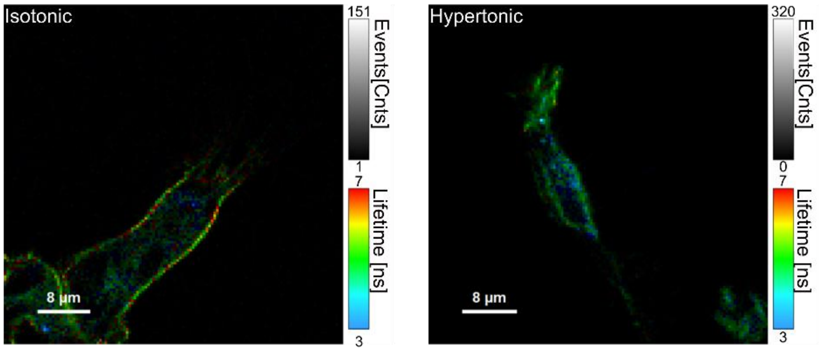


**Figure S2**. Change in membrane tension under hypertonic conditions in differentiated HL-60 cells (dHL-60). dHL-60 show differences in membrane tension under hypertonic conditions compared to isotonic conditions.

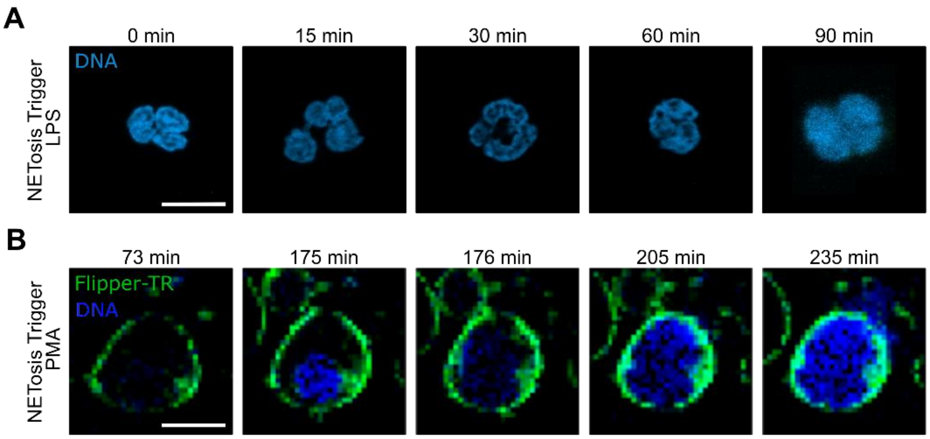


**Figure S 3**. A) Chromatin staining (DAPI) of neutrophilic granulocytes during LPS induced NETosis. Scale bar = 10 µm. B) False-color representations of membrane (Flipper-TR, green) and nucleus (SPY505, blue) staining during the process of PMA induced NETosis in neutrophilic granulocytes (intensity-based images). Scale bar = 10 µm.

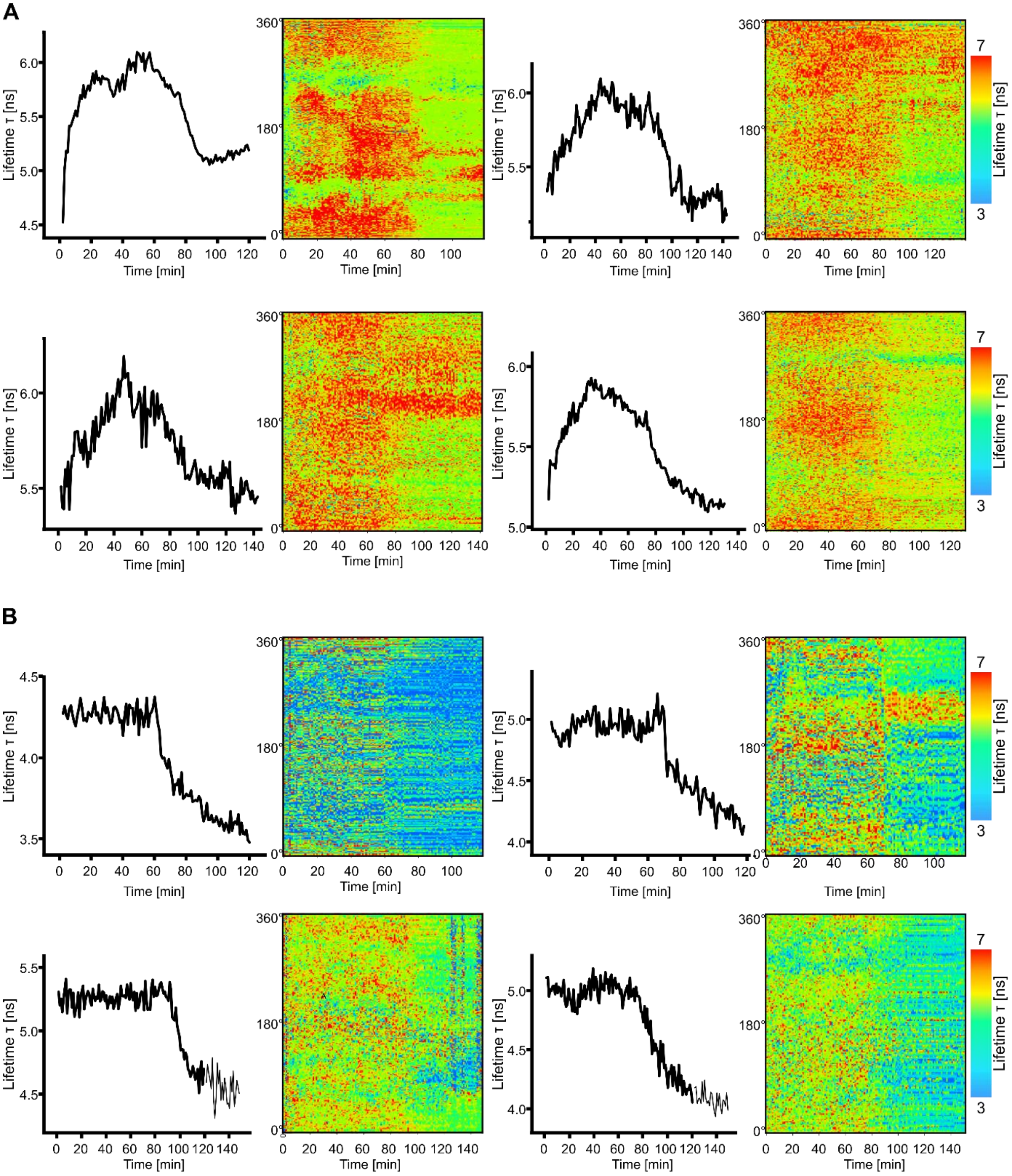


**Figure S4**. A) Mean fluorescence lifetime (τ) for exemplary control neutrophils with their corresponding heat map visualization of membrane lifetime (τ) values at each time point - from PMA trigger (-20 min) to post-rupture. This representation provides a spatial and temporal overview of membrane tension dynamics. B) Mean fluorescence lifetime (τ) for exemplary cholesterol depleted neutrophils with their corresponding heat map visualization of membrane lifetime (τ) values at each time point - from PMA trigger (-20 min) to post-rupture.

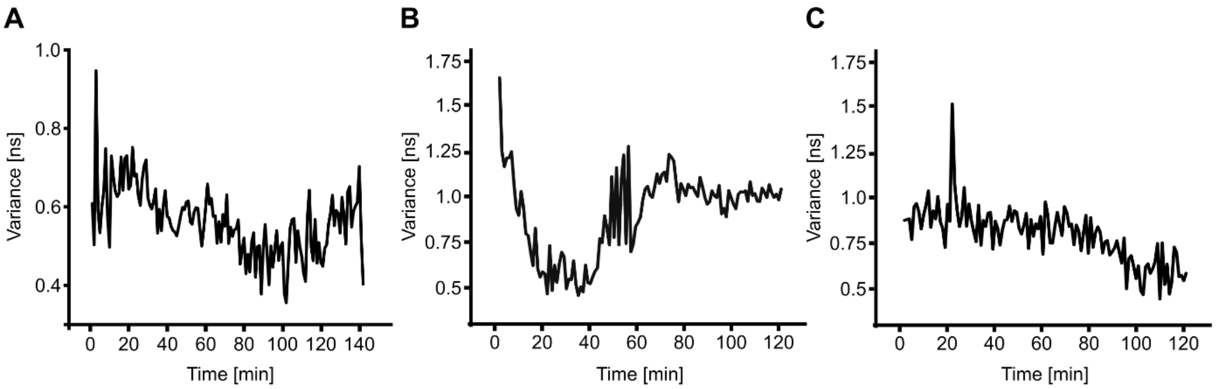


**Figure S 4**. Variance of membrane tension. A) For the exemplary neutrophil shown in Figure 2D, we calculated the variance of lifetime values derived from each pixel intensity along the cell contour ROI (membrane) at every time point, showing fluctuations in membrane tension over time. B) Variance of lifetime values at each time point for the exemplary control neutrophil illustrated in Figure 3C, which also displays noticeable temporal fluctuations and a less uniform pattern. C) Variance of lifetime values at each time point for the exemplary cholesterol-depleted neutrophil illustrated in Figure 3C, which appears comparatively more homogeneous over time, indicating reduced variability.

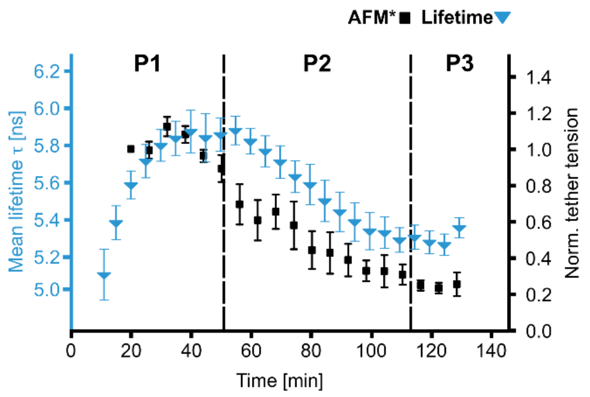


**Figure S 6**. Combined lifetime data (cell membrane) with previously reported[1] AFM data showing cell stiffness (Young's modulus) of life neutrophilic granulocytes after stimulation with PMA. Both lifetime and stiffness increase after PMA stimulation and decrease afterwards again. n(lifetime) = 9, n(AFM) = 3. Mean ± SEM

### Material and Methods

Neutrophil granulocytes isolation

For the isolation of human neutrophil granulocytes, the EasySep™ Direct Human Neutrophil Isolation Kit (Stemcell technologies, Vancouver; Canada) is used. In this process, 7.5 ml of human blood is collected into a sterile K3 EDTA S-Monovette® (Sarstedt, Nümbrecht, Germany) and allowed to cool to room temperature (RT). Next, 3-4 ml of the blood is transferred to a 15 ml Falcon tube (Sarstedt, Nümbrecht, Germany). To this, 50 µl/ml of isolation cocktail and 50 µl/ml of rapid spheres™ are added, followed by a 5-minute incubation at RT.

The sample is then brought to a total volume of 10 ml using PBS with 1 mM EDTA, inverted for mixing, and placed in a ring-shaped magnet for 5 minutes. The supernatant is transferred to a new Falcon tube with the same amount of rapid spheres™ added again, followed by another 5-minute incubation outside and then inside the ring-shaped magnet.

This process is repeated once more, with the supernatant being transferred to a new Falcon tube. The isolated cells are then centrifuged for 3 minutes at 60 g, the supernatant is discarded, and the cell pellet is carefully resuspended in 1 ml of clear Roswell Park Memorial Institute 1640 medium (clear RPMI 1640) for further use.

Cell culture

HL-60 cells used in the experiments have been purchased from German Collection of Microorganisms and Cell Cultures GmbH (DSMZ ACC 3) and were maintained in RPMI 1640 medium (Gibco) supplemented with 10% Fetal Bovine Serum (Gibco) and 1% penicillin/streptomycin (Gibco) at 37 °C and 5% $CO_2$. The cells were passaged every second to third day, keeping the density between $10^5$ and $2*10^6$ cells/ml. All-trans retinoic acid (ATRA) (Sigma Aldrich, St. Louis, USA) and DMSO (Sigma Aldrich, St. Louis, USA) were used to promote neutrophil-like differentiation of HL-60 cells. A stock solution of 10 mM ATRA was prepared in dimethysulfoxide (DMSO) and stored at -20 °C in the dark. Cells were seeded at 300,000 cells/ml in a 3.5 cm glass bottom cell culture dish (MatTek, Ashland, USA) containing the cultivation medium with an additional 4 µM ATRA and 1% DMSO. The medium was changed every 2-3 days.

HeLa cells were purchased from DSMZ (DSMZ ACC 57), cultured at 37 °C and 5% $CO_2$ in IMDM (Gibco) with 1% Pen/Strep (Gibco) and 10% fetal bovine serum (Gibco). Cells at 80-90% confluence were trypsinated and split 1.10 every 3-4 days. For measurements, $3*10^4$ cells were seeded in a 3.5 cm glass bottom dish (Bioptechs, PA, USA) and cultured for two days. HeLa cells were washed with DPBS and stained with 1 µM Flipper-TR in growth medium for 20 min at 37 °C and 5% $CO_2$. Afterwards, the staining solution was removed and exchanged to either hypertone medium (1.6 M sucrose solution) or hypotone medium ($ddH_2O$).

Flipper-TR Labeling of Cells

Flipper-TR was dissolved in DMSO according to the manufacturer's instructions. HeLa and dHL-60 cells were labelled with 1 µM Flipper-TR (Spirochrome AG, Stein am Rhein, Switzerland) in growth medium and neutrophilic granulocytes were labelled with 250 nM Flipper-TR in clear RPMI medium without supplements for 20 min at 37 °C and 5% $CO_2$. The staining solution was not removed prior to imaging, as the dye remained membrane-associated.

For experiments including phorbol-12-myristat-13-acetat (PMA, Merck) stimulation, PMA was added to reach a final concentration of 100 nM and cells were incubated for 10–15 min at room temperature before imaging.

Dual Labelling With SPY505 and Flipper-TR

SPY505 (Spirochrome AG, Stein am Rhein, Switzerland) was prepared in DMSO as per the manufacturer's recommendations. Cells were first incubated with 2 µL SPY505 diluted in RPMI medium for 40 minutes at 37 °C, 5% $CO_2$. Subsequently, Flipper-TR diluted in RPMI was added to a final concentration of 250 nM and cells were incubated for another 20 minutes under the same conditions. PMA stimulation was performed as described above.

Cholesterol Depletion

Cholesterol depletion was performed using methyl-β-cyclodextrin (MβCD, Sigma Aldrich, St. Louis, USA). A stock solution was prepared in water. Cells were incubated with 5 mM MβCD in RPMI medium for 1 hour. After treatment, cells were labelled sequentially with SPY505 (40 minutes), Flipper-TR (20 minutes), and stimulated with 100 nM PMA prior to imaging.

Microscopy Settings

Experiments were carried out on a MicroTime 200 system (PicoQuant) using a 60× water-immersion objective. Excitation was performed at 481 nm. Detection was split onto two SPAD detectors equipped with appropriate filters to separately record Flipper TR (594 LP) and SPY505 (530/20) emission. Images were acquired in accurate scanning mode at 128 × 128 pixels, with a dwell time of 2 ms per pixel (scan time 50 s per image).

NETosis with LPS

After isolation neutrophils were stimulated with 10 µg/mL LPS for the indicated time points. The cells were then fixed with 4% paraformaldehyde for 15 min. The cells were then stained with Hoechst for nuclear staining for 10 min (1:10000).

Data analysis

Lifetime data were evaluated using the PicoQuant software and a python code. The Code is available upon request.

Fluorescence lifetime–based membrane tension analysis

Fluorescence lifetime imaging data were processed using a custom-written Python pipeline based on NumPy, SciPy, OpenCV, Pandas, and Matplotlib. For each experiment, time-resolved intensity data were imported from text files, while corresponding pseudo-colored lifetime images were loaded from bitmap (BMP) exports. Lifetime values were reconstructed from RGB-encoded images by normalizing individual color channels to total intensity and calculating lifetime values using empirically determined linear channel weights.

Cell segmentation and contour detection

Cells were identified in the first time frame based on intensity thresholding. Regions of interest were defined either automatically by selecting the largest contiguous objects or manually using user-defined coordinates. For each cell, a bounding box was applied consistently across all time points. Cell contours were detected by adaptive thresholding of grayscale intensity images followed by morphological smoothing. To avoid edge artifacts, contours were eroded by a defined pixel offset, yielding an inner membrane contour used for lifetime readout.

Spatial sampling of membrane lifetime

For each time point, membrane pixels were extracted along the detected contour. Contour coordinates were ordered angularly around the cell centroid to ensure consistent spatial orientation across time. Fluorescence lifetime values were sampled along the membrane and stored as one-dimensional profiles representing spatial membrane tension distributions. For visualization and comparison across time, profiles were interpolated to a common length and displayed as heat maps.

Temporal analysis

Mean membrane lifetime values were calculated for each cell and time point. Lifetime values were clipped to a defined range to reduce noise. Temporal evolution of mean lifetime was used as a proxy for changes in membrane tension over time. All intermediate data, including contour coordinates, thresholds, and lifetime values, were stored in CSV and JSON formats to ensure reproducibility.
